# A Novel Approach to T-Cell Receptor Beta Chain (TCRB) Repertoire Encoding Using Lossless String Compression

**DOI:** 10.1101/2023.01.30.526195

**Authors:** Thomas Konstantinovsky, Gur Yaari

**Affiliations:** Faculty of Engineering, Bar Ilan University, 5290002 Ramat Gan, Israel; Bar Ilan Institute of Nanotechnology and Advanced Materials, Bar Ilan University, 5290002 Ramat Gan, Israel

## Abstract

T-cell diversity is crucial for producing effective receptors that can recognize the pathogens encountered throughout life. A stochastic biological process known as VDJ recombination accounts for the high diversity of these receptors, making their analysis challenging. We present a new approach to sequence encoding and analysis, based on the Lempel-Ziv 76 algorithm (LZ-76). By creating a graph-like model, we identify specific sequence features and produce a new encoding approach to an individual’s repertoire. We demonstrate that this repertoire representation allows for various applications, such as generation probability inference, informative feature vector derivation, sequence generation, and a new measure for diversity estimation.

## Introduction

Living in an environment where pathogens are a constant threat, our adaptive immune system has little to no time to rest, constantly trying to recognize pathogens by binding T- and B-cell specific receptors to their antigens. In order to produce a proper receptor that could bind to a specific antigen, our body leverages the stochastic V(D)J recombination process, which accounts for the high receptor variability seen in individuals [17]. Understanding the fine details of the mechanism behind the V(D)J recombination and how they affect the dynamics and variability of our adaptive immune system is one of the fundamental open questions in immunology. A proper methodology for a compact representation of an entire repertoire, which encapsulates as much information as possible, is of crucial need. Such a representation will provide a potent and convenient way to analyze and infer the different attributes that compose an individual repertoire. While it is a well-established fact that the V(D)J recombination process is highly non-deterministic [9, 17, 14], numerous models were suggested to capture recombination statistical properties [32, 21, 31, 24, 30]. Few methods were proposed aiming to convert the large number of sequences in an individual repertoire into a single vector representation [27, 29, 38, 8, 26, 36]. Many applications revolving around the storage and processing of data produced by high-throughput processes leverage various compression algorithms to apply relevant producers in an efficient manner. These compression algorithms are implemented via different methodologies offering a spectrum of compressed formats and are commonly separated into two groups. A compression algorithm is defined as either lossless (i.e, when a compressed file is decompressed, the output matches the original file) or lossy (i.e., when a compressed file is decompressed, the output is epsilon-close to the original data, but not identical). Standard lossless compression algorithms that are being used in many domains [5, 4, 1, 37, 15] are based on deriving a token-based mapping to reduce the compressed file size [16, 18, 7, 35, 10, 3, 13]. Yet, there are no published studies attempting to combine these concepts from information theory, explicitly leveraging the various lossless data compression algorithms as feature extractors. A related interesting question is whether such a compressed representation of a repertoire, alongside the token dictionary used to generate it, can lead to a statistical model that captures the inner workings of a repertoire. Here we propose a new methodology for encoding the CDR3 regions of the T-cell receptor beta chain repertoire into a single graph-based representation, utilizing the Lempel-Ziv compression algorithm (LZ-76) [18]. The LZ-76 algorithm is a lossless data compression technique that uses a sliding window to look for repeated patterns in input data. It replaces the repeated patterns with references to previous occurrences in the data and thus results in a compressed output. The encoding approach we propose allows for in-depth repertoire analysis, feature extraction for statistical modeling, synthetic sequence generation, generation probability inference, and a new measure for diversity estimation. We elaborate and demonstrate how, given a repertoire, the resulting model captures its inner dynamics, allowing us not only to classify cohorts of individuals based on their repertoires, but also to generate new synthetic sequences and infer generation probability (*P*_*gen*_) for each sequence. The resulting repertoire graph model aims to provide researchers with a new analytic perspective on adaptive immune receptor repertoires (AIRRs), one that originates from the nucleotide/amino acid sequence structure only.

## Methods

### Data

The datasets used in this work are composed of TCR beta chain CDR3 sequences. They are summarized in table 1.

**Table 1:** A summary of the datasets used throughout this paper.

| Dataset Abbreviation | Condition | Sample Size | Reference |
| --- | --- | --- | --- |
| DS1 | HIV | 192 | [34] |
| DS2 | HIV | 39 | [33] |
| DS3 | CMV and controls | 785 | [11] |
| DS4 | COVID | 950 | [25] |

The development and analysis of the methods were performed on productive CDR3 sequences in both nucleotide and amino acid scopes. TRBV and TRBJ gene annotations were taken for each sequence as provided via “ImmunoSeq Access” [6]. No further preprocessing was applied to the repertoire data.

### LZ-76

The Lempel-Ziv dictionary is defined as the number of different sub-patterns encountered as each sequence is viewed from beginning to end. The process of sequence tokenization was achieved by applying the most basic form of the Lempel-Ziv algorithm [39, 18] to derive a dictionary of unique sub-patterns (Figure 2). Since there is no natural order of the CDR3 sequences in a repertoire, a dictionary of sub-patterns was derived per sequence, thus resulting in *N* dictionaries where *N* is equal to the input repertoire depth.

**Figure 1:**
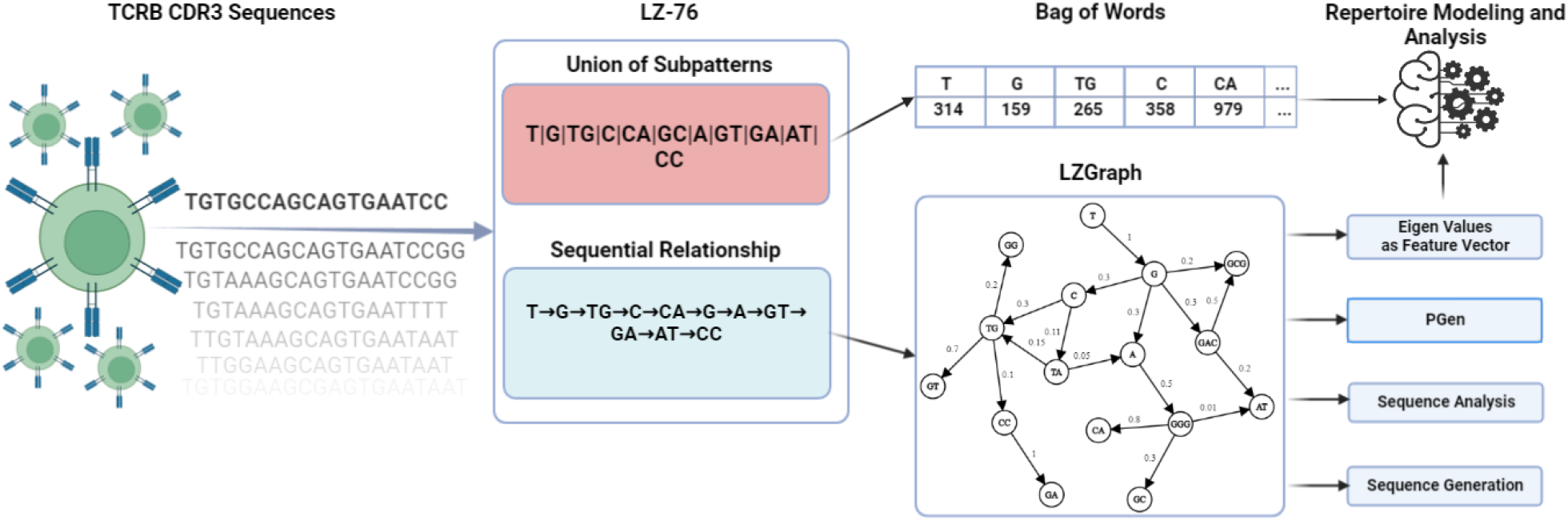
An illustration of the proposed methodology flow, from sequence to model: from right to left the transition between T-cell receptors and their corresponding CDR3 sequence, to the various models and applications on the right.

**Figure 2:**
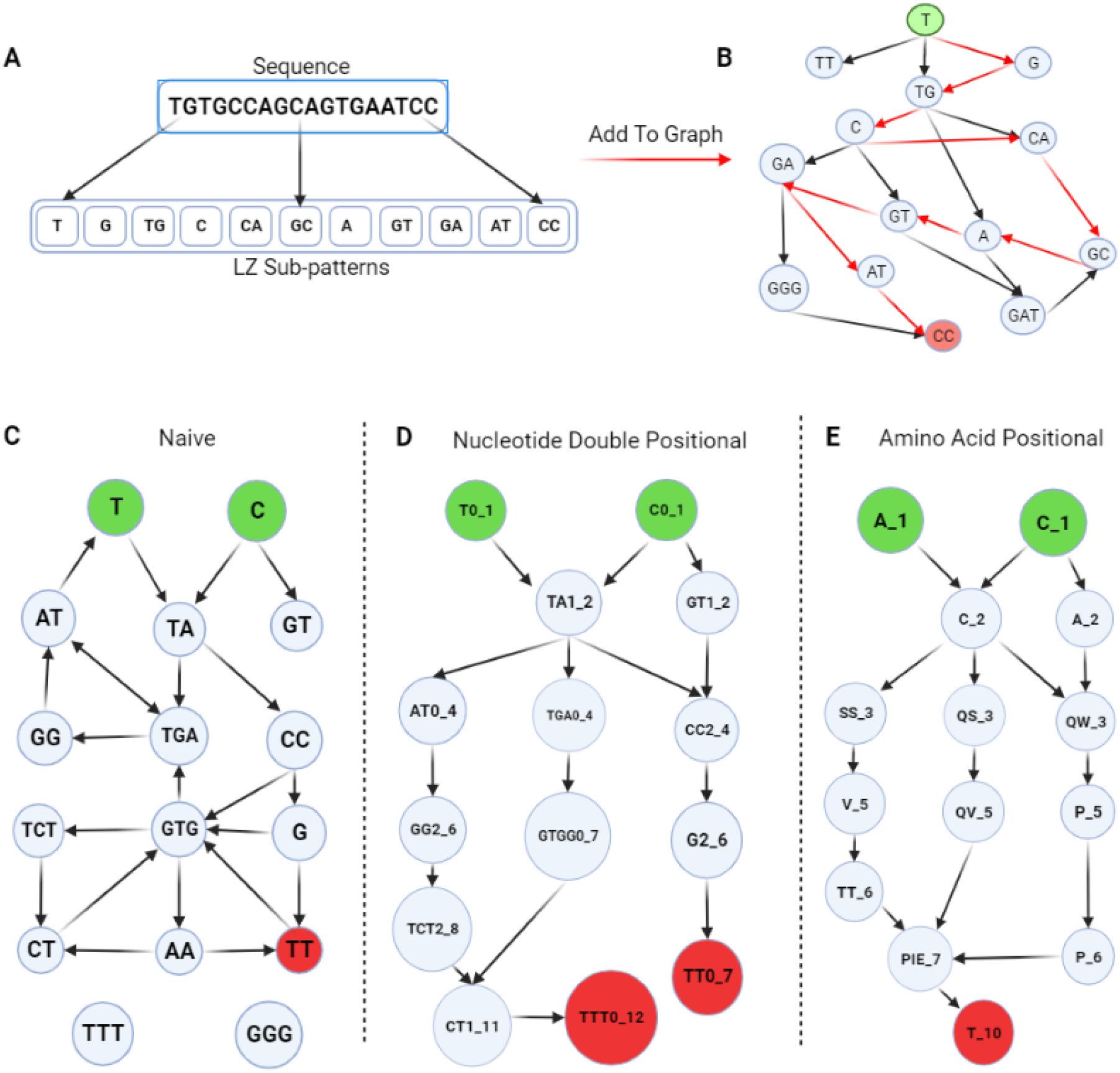
A summary of sequence to graph transitions. (**A**) is an illustration of LZ-76 decomposition of a sequence into sub-patterns; (**B**) is how the LZ-76 sub-patterns extracted in (**A**), integrate into a graph structure. Green nods are graph initial states, and red nodes are terminal states. The red arrows represent the newly added/updated edges extracted from the sequence in (**A**). (**C**) is an example of the “Naive” graph structure not respecting the DAG constraint. **(D)** is an example of the “Nucleotide Double Positional” graph, respecting the DAG constraint, differentiating nodes by the reading frame start position and the position in a sequence. **(E)** is an example of the “Amino Acid Positional” graph, respecting the DAG constraint, differentiating nodes by the reading position in the sequence.

A common way to understand the use case of LZ-76 is by considering it as an approximation method for estimating the value of the theoretical Kolmogorov complexity, which is the length of the shortest computer program that is able to reproduce a given pattern [19]. LZ-76 is also an efficient estimator of the entropy rate, which measures how many bits of innovation are introduced by each observation [2]. Naturally, the more complex a sequence is, the higher the entropy, the more resources will be needed to recreate that sequence, i.e., a higher Kolmogorov complexity.

Due to the greedy nature of LZ-76 algorithm, each time a new sub-pattern is observed it is added to the aggregated vocabulary. A special case has to be addressed when the last observed sequence of symbols is already a sub-pattern in our dictionary. Our solution consists of appending those symbols to the last observed sub-pattern.

### LZ-76 Bag of Words Representation

One of the approaches that we propose for repertoire representation is a bag of words (BOW): a frequency table of the set of unique sub-patterns from all sequences observed in a repertoire. This frequency table is referred to as BOW feature vector. The resulting representation encapsulates the overall repertoire composition in terms of LZ-76 sub-patterns. However, an important piece of information is discarded, the order of the sub-patterns. To capture both the sub-pattern structure of a repertoire and the sequential relationships between those sub-patterns, a graph data structure was leveraged.

### Graph-Based Representation (LZGraph)

The nodes of an LZGraph are the set of unique sub-patterns. The edges of the graph are formed by connecting sequential sub-patterns, as the sequences are being processed one by one. The weights assigned to each edge in the graph are proportional to the empiric transition events between the connected nodes, normalized by the out-degree of each node. An additional layer of information can be added to each edge documenting the relative V and J gene usage of the sequences contributing to this edge.

Formally the LZGraph of repertoire *x* is defined as the set triplet (*V, E, W*) that stands for vertices (aka nodes), edges, and weights, respectively.

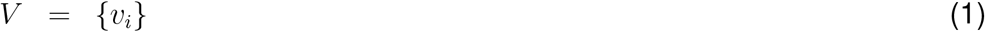

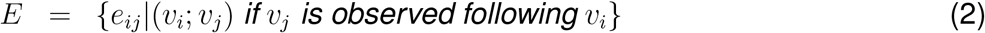

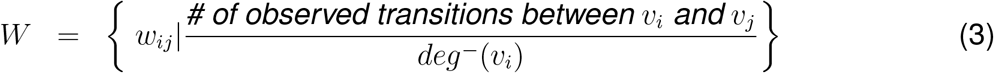

∀*i* 1..*N*. *deg*^−^(*v*_*i*_) indicates the out-degree of node *v*_*i*_. Two additional sets are the set of all initial states (*Init*), and the set of all terminal states (*Term*).

We present three main graph types: “Naive”, “Nucleotide Double Positional”, and “Amino Acid Positional” graphs. “Naive” refers to a graph whose nodes have no additional positional information (each node is an LZ-76 sub-pattern only). The nodes of the “Nucleotide Double Positional” graph appends two additional pieces of information to the “Naive” node representation. On top of the LZ-76 sub-pattern, they include the reading frame start position and the absolute location in the sequence. In general, the form of each node is defined as {LZ-76 sub-pattern} {modulo 3 position (reading frame)} {absolute position}. Lastly, the nodes of the “Amino Acid Positional” graph are the LZ-76 sub-patterns of the CDR3 amino acid (aa) sequence, appended by the start position of the sub-pattern in the sequence. For example, CA 2 would be a node in the graph, which is interpreted as the sub-pattern “CA” (Cysteine-Alanine), starting at the second position in the CDR3 aa sequence.

The insertion of positional components to each sub-pattern in the non-naive graphs enforces the resulting LZGraph to be a directed a-cyclical graph (DAG). A DAG graph ensures that there is a constant flow from the root node (a node belonging to the *Init* set defined above) to the terminal node (a node belonging to the *Term* set defined above), avoiding any self-loops or cycles which in a “Naive” graph occur naturally. Adding positional information allows for greater interoperability of the graph, as each node can be associated with a specific position in the CDR3 sequence.

### Generation Probability (*P*_*gen*_)

*P*_*gen*_ is a measure to estimate the generation probability of a given sequence. It can be calculated using different generation models. LZ-76 *P*_*gen*_s are derived from the graph types presented above. The interpretation of these *P*_*gen*_s depends on the graph type used.

The LZGraph *P*_*gen*_ of a given CDR3 Amino Acid/Nucleotide sequence s is formally defined as:

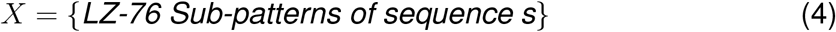

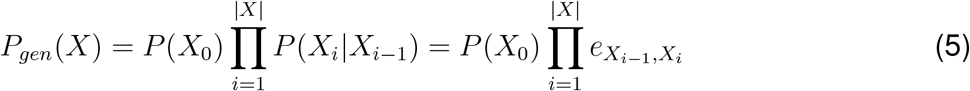

where *P* (*X*_0_) is the marginal probability of *X*_0_ ∈ *Init*, and *X*_|*X*|_ ∈*Term*.

In a scenario where edges exist in a given sequence but were not observed while constructing the LZGraph, the missing values are imputed with the geometric mean of the non-missing edges of the same sequence.

### Graph Based Feature Vector

There are multiple ways to extract a vector representation of a graph. Here we used the eigenvector centrality as the feature vector to describe a repertoire. It can be interpreted as the extent to which nodes influence the propagation in the network. The eigenvector centrality (*x*) of a graph is calculated by solving the following equation:

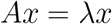

where *A* is the adjacency matrix of the graph and *λ* is the largest eigenvalue of *A*. The eigenvector centrality of node *v*_*i*_ ∈ *V* is the *i*’th component of *x*, namely *x*_*i*_. A node with a high eigenvector centrality value is characterized by being adjacent to other nodes with high eigenvector centrality values.

### Sequence Analysis

We propose two new approaches for sequence analysis that can be used for both gene annotation inference of a sequence and a comparison between sequences.

1. Sequence Variation Curve The sequence variation curve (Figure 7A) represents the number of possible immediate alternatives for each LZ-76 sub-pattern of a given sequence. Given a sequence *S*, it is first decomposed into its LZ-76-sub-patterns denoted as *Ŝ* The curve *C* is then defined as:

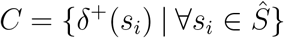

where *δ*^+^(*s*_*i*_) is the number of child nodes from node *s*_*i*_ based on a given LZGraph.
2. Gene Variation Plot

The gene variation plots (Figure 7B/C/D) depict the number of different V and J genes that are associated with a certain node/edge for a given sequence *S*. It is first decomposed into its LZ-76-sub-patterns denoted as *Ŝ*, for each pair of sequential subpatterns in *Ŝ* we extract the observed alleles that are associated with such a transition during the construction of the LZGraph. Alleles that are absent in more than *alpha* percent of the sequence path are omitted; the names of the alleles that have been observed through the entire path are colored in red. The process is repeated for both V and J genes separately. The resulting overview (Figure 7C/D) shows the likelihoods of different alleles for each edge along the sequence path.

### Sequence Generation

LZGraph can be used to simulate new sequences with similar statistical properties to the source sequences that were used to construct the graph. Here we applied two approaches for simulating new sequences, “Unconstrained” and “Genomically Constrained”

1. Unconstrained Method The Unconstrained method for sequence generation randomly selects an initial state from the *Init* set based on the empirical probabilities. To the initial sub-pattern more nodes are sequentially appended in a stochastic fashion, where the next node is drawn based on the weight distribution of the emanating edges. When a node from the *Term* set is encountered, a conditional stopping procedure is carried out. The conditional stopping procedure checks whether there are any other nodes from the *Term* set that could be reached from the current node. In the case that there are no such nodes, the algorithm halts and uses the encountered node as its terminal state and as the final piece of the generated sequence. On the other hand, if the above does not hold, all the other nodes from *Term* that are reachable from the current node are used to derive a stopping probability *P*_*T*_. *P*_*T*_ (*T*_*i*_) is defined as the probability of stopping at a given terminal node (*T*_*i*_∈ *Term*) conditioned on not stopping at any other terminal state that can be reached from *T*_*i*_. Formally defined: Let Ψ(*T*_*i*_) be the probability of stopping at *T*_*i*_ regardless of the initial conditions. I.e., Ψ(*T*_*i*_) equals to the number of paths terminated at *T*_*i*_ divided by the total number of paths used to construct the graph. Hence

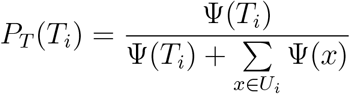

where

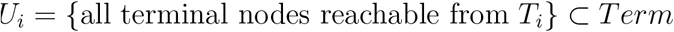
2. Genomically Constrained

In the Genomically Constrained method, alongside a random initial state, a random V gene and a random J gene are selected based on the relative frequency observed at the LZGraph’s source repertoire. Similar to the unconstrained method, as long as the generation process of a sequence did not reach a terminal state, the next node is selected randomly based on the relative weight distribution of the out-edges of the current node. Since these out-edges can be associated to any V and J genes, in this method we take into account only edges that have non zero support for these selected genes. In other words, when building the LZGraph, if we never observed a sequence having the V and J genes that had a transition between the current node and the next node candidate, this edge is excluded from the possible next step.

### Running Sonia

The comparison with Sonia was performed using the python open-source library sonia version 0.1.2. The “SoniaLeftposRightpos” model was fitted using the “human T beta” chain type, and the repertoire of choice as the “data seqs” argument of the model. 200K sequences were generated from the fitted model and were used to further fine-tune it using the “add generated seqs” method followed by 30 epochs of “infer selection” converging on the best fit. For the generated sequence comparison, the library’s “SequenceGeneration” wrapper was used to generate post sequences via the “generate sequences post” method. For *P*_*gen*_ comparison the library’s “EvaluateModel” wrapper was used to evaluate a set of sequences, and the “ppost” result was used for the comparisons.

## Results

### LZ-76 Bag-of-Words(BOW) Vectors to Encode Repertoires

#### The LZ-76 BOW Dictionary Generated From a Repertoire Encapsulates Valuable Information

The dictionary generated from applying LZ-76 to a repertoire can be used as a BOW feature vector to represent it. To assess how well such a representation encapsulates the unique patterns of different conditions, we performed the following experiment: For each dataset, we first used all relevant repertoires (Table 11) to create a dictionary of unique LZ-76 sub-patterns, aka the BOW dictionary. All repertoires were then encoded using the pre-calculated dataset specific LZ-76 BOW dictionary. The counts for each repertoire BOW vector were then normalized by the repertoire depth. The resulting encoded repertoires (BOW vectors) were then projected onto R^2^ using UMAP [22]. The same procedure was carried out for each dataset in Table 1. The collection of repertoire projections was then examined for any noticeable spatial patterns in the embedded 2D space (Figure 3). Although dictionary sizes and the distribution of frequencies over the LZ-76 sub-patterns slightly vary between the three datasets (see Figure 3B), the overall projection in all three cases is robust to the choice of dictionary. A projection of the repertoires from all datasets is shown in Figure 3A, where the LZ-76 BOW vector is constructed using the dictionary based on sub-patterns observed in DS1+DS2. Projections using dictionaries from the other datasets are shown in supplementary figure-1. Even though there are sub-patterns unique to DS3 and DS4 that do not appear in DS1+DS2 (see Figure 3), by encoding the repertoires using the DS1+DS2 dictionary only, we observe a clear segregation between the sets of projected repertoires (most likely due to batch effects; Figure 3). Furthermore, if we consider that DS4 is composed of a few smaller datasets, it is interesting to see that, nevertheless, their mapping into the 2D space is similar (see supplementary figure-1.). Alongside projecting the data onto R^2^, we trained a simple decision tree classifier to validate the batch effect hypothesis. We used three different dictionaries derived for each one of the datasets, DS1+DS2, DS3, and DS4. We classified the different sets using such a model, with an average micro F1 score of 0.89 ± 0.02 on 5 folds of cross-validation.

**Figure 3:**
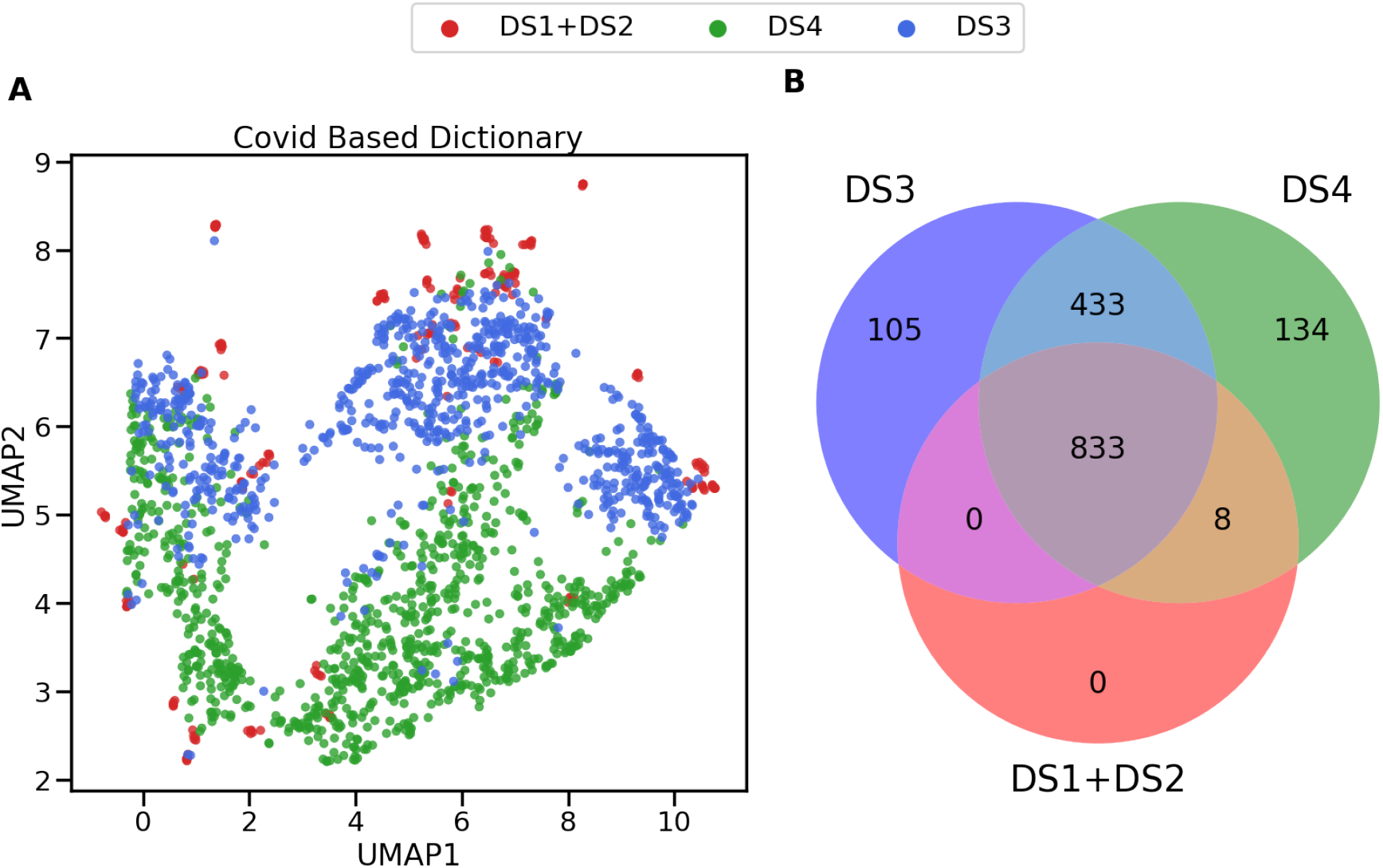
BOW dictionary encoded repertoires spatial clustering and dictionary similarity. (**A**) All four datasets were projected onto ℝ^2^ using UMAP. The features used to project each repertoire were all sub-patterns discovered in DS4. Each color represents a dataset. (**B**) Venn Diagram of unique LZ-76 sub-patterns observed in each dataset.

### LZ-76 BOW Vectors Are Robust Features For Classification Tasks

After showing that LZ-76 BOW vectors can be used to identify phenomena such as batch effects between datasets, we tested the applicability of this approach for classifying females and males, in four distinct datasets. We first compared LZ-76 BOW feature vectors to K-mer based vectors. For this, we used DS1 to train an AdaBoost model and compared the resulting F1 scores via 15 folds of cross-validation to rule out any batch effect (see Figure 4A). The robustness of the BOW feature vectors was then evaluated by introducing a second dataset to the already trained model, namely DS2, while ignoring any new sub-patterns that might have appeared while encoding it. Along with surpassing the traditional K-mer approach, the LZ-76 BOW feature vector allowed the trained model to be more robust, as demonstrated by its significantly higher F1 score (dashed red line in Figure 4A).

**Figure 4:**
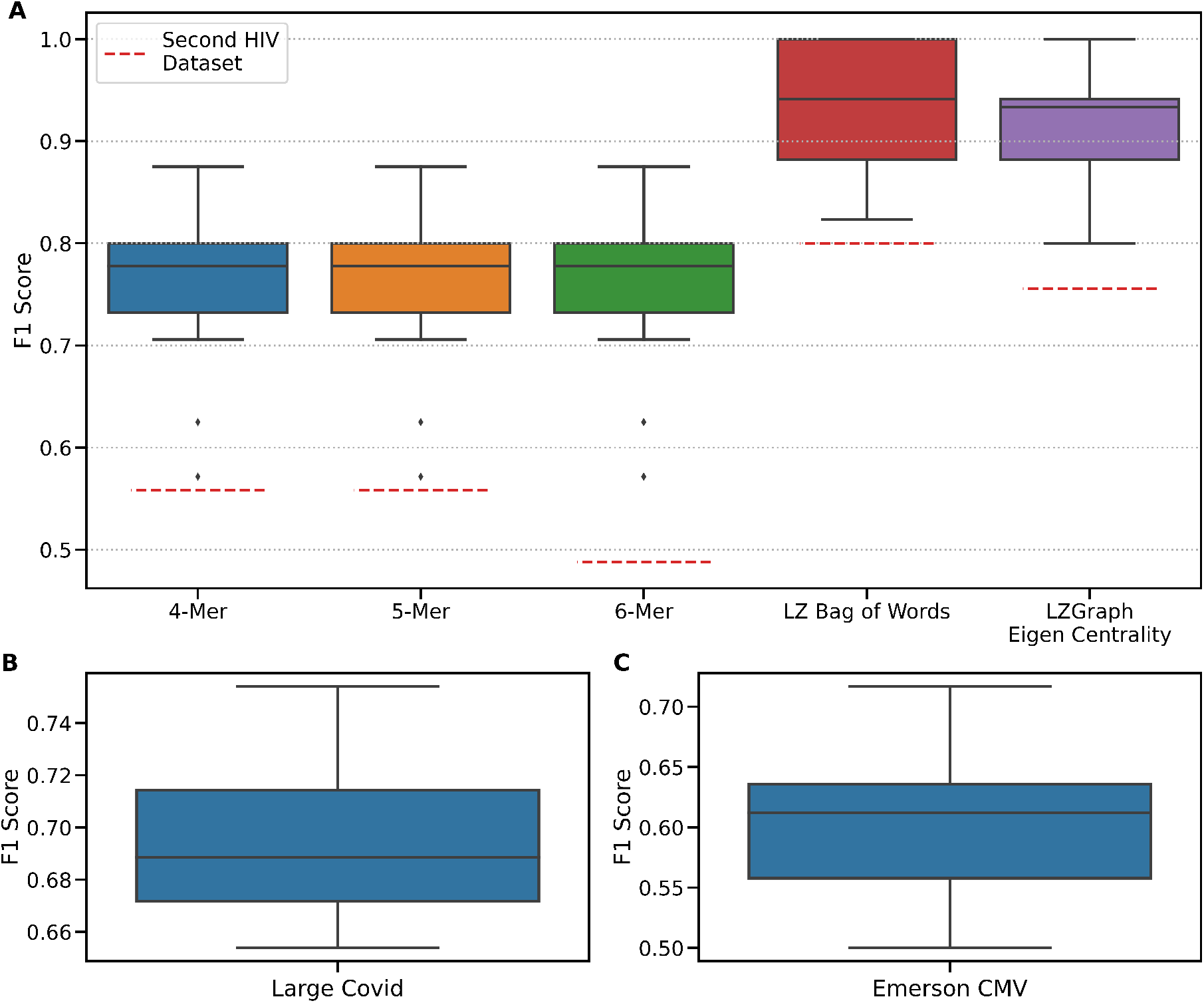
The Resulting F1-Scores of Multiple Predictions Made Using LZ-76 and K-mer Features. F1 scores of a 15-fold cross-validation on an AdaBoost model predicting the sample’s sex. (**A**) features derived from 5 different extraction methods are represented by the colored boxes. The dashed red line represents the F1 score resulting from the prediction on DS2, using the model trained on DS1. (**B**) F1 scores of 15-fold cross-validation on a test set of 300 samples out of 950 samples from DS4. (**C**) F1 Scores of 15-fold cross-validation on a test set of 230 samples out of 785 samples from DS3.

### Encapsulating Sequential Relationships via “LZGraphs” Results in an “all-in-one” Sequence Analysis Model

Although we observed sufficient information encapsulation via the “naive” LZ-76 BOW representation, an important piece of information is lacking in such a setting, namely, the sequential relationships between the LZ-76 sub-patterns. The naive setting does not impose any constraint on the sub-patterns, in fact, the naive setting is no more than normalized count vectors (BOW). By transitioning to a graph (LZGraph), we reattain the sequential relationship between the sub-patterns as well as the probabilities associated with those transitions. In this paper, we tested three versions of an LZGraph: Naive Nucleotide, Nucleotide double positional, and aa positional (see Figure 2 in the methods section). Such representations open the door for a vast variety of applications, some of which are presented below.

#### The number of nodes and edges in an LZGraph scale as a Power-law with the number of its’ unique source sequences

The number of unique TCRs in the human body is enormous (∼10^10−11^; [20]). Nevertheless, it constitutes only a tiny fraction of the space of theoretically possible receptors. If we assume that the average CDR3 aa length is ∼15 [28] and ignore the finite number of deterministic patterns that can appear at the beginning and end of the sequence (resulting from the V(D)J recombination process), the theoretical maximum number of sequences would be 20^15^ ∼3.3· 10^19^.

To measure the connection between the size of an LZGraph and the number of empirical sequences that were used to construct it, we took the following approach. For each dataset we incrementally added repertoires and constructed from the accumulated set of sequences a “Nucleotide Double Positional” LZGraph (see methods). We repeated the above procedure 10 times, shuffling the order of added repertoires. Figure 5 shows the relationships between the number of sequences used for the construction of the graph for DS1+DS2, DS3 and DS4 and the number of nodes (panels A and B. Edges are shown in D and E). It is clear that even for 785 repertoires (DS3) containing 150M sequences, the two curves are not saturating (Figure 5A-B). Plotting these relations in a log-log plot, we observe power-law dependencies. In particular, the number of nodes scales with the number of sequences with a power-law exponent of 0.25-0.28, depending on the dataset, and the number of edges with an exponent of 0.43-0.51, depending on the dataset (Table 2). The differences between the datasets may be linked to differences in the diversities in these datasets, but in a non-straightforward way as the respective Hill diversity curves [12] of the samples in these datasets are entangled. These power-law relationships demonstrate that individual repertoires carry a wealth of unique patterns that can add new information, even in cases of graphs that were built from thousands of repertories.

**Table 2:**
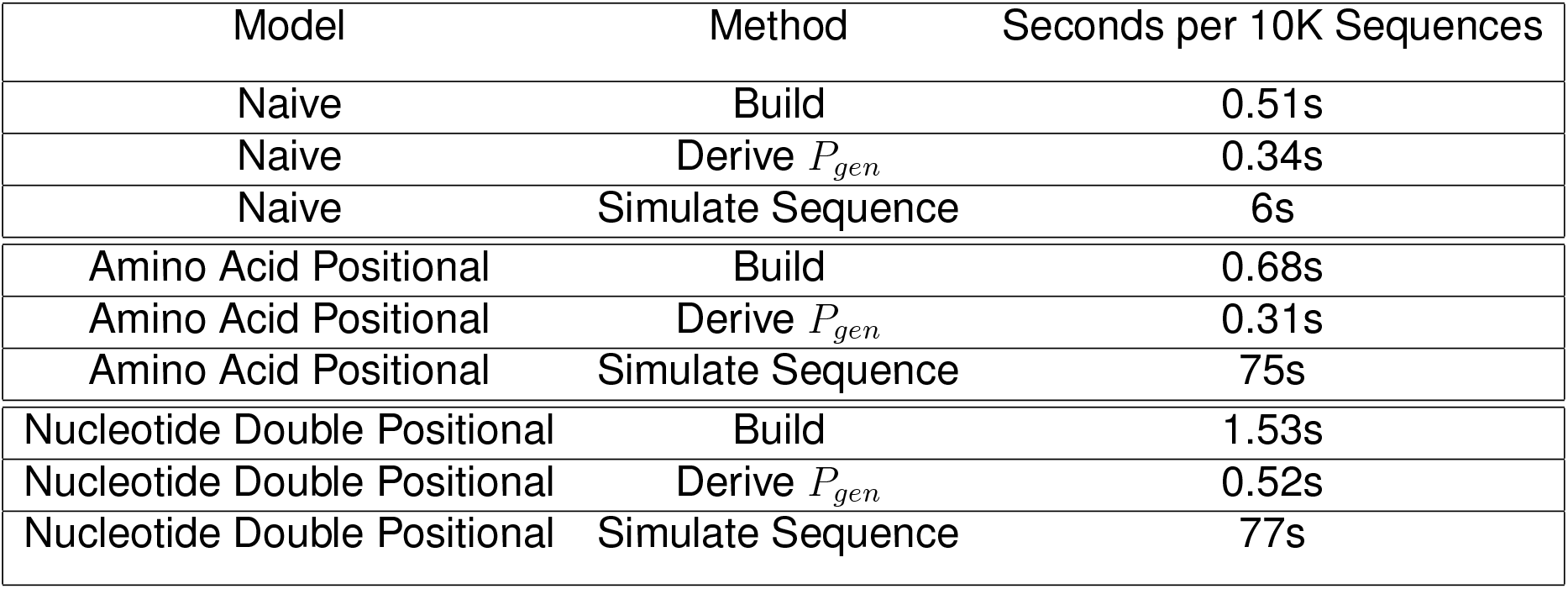
A summary of run-times of the different models and methods presented in this paper.

**Figure 5:**
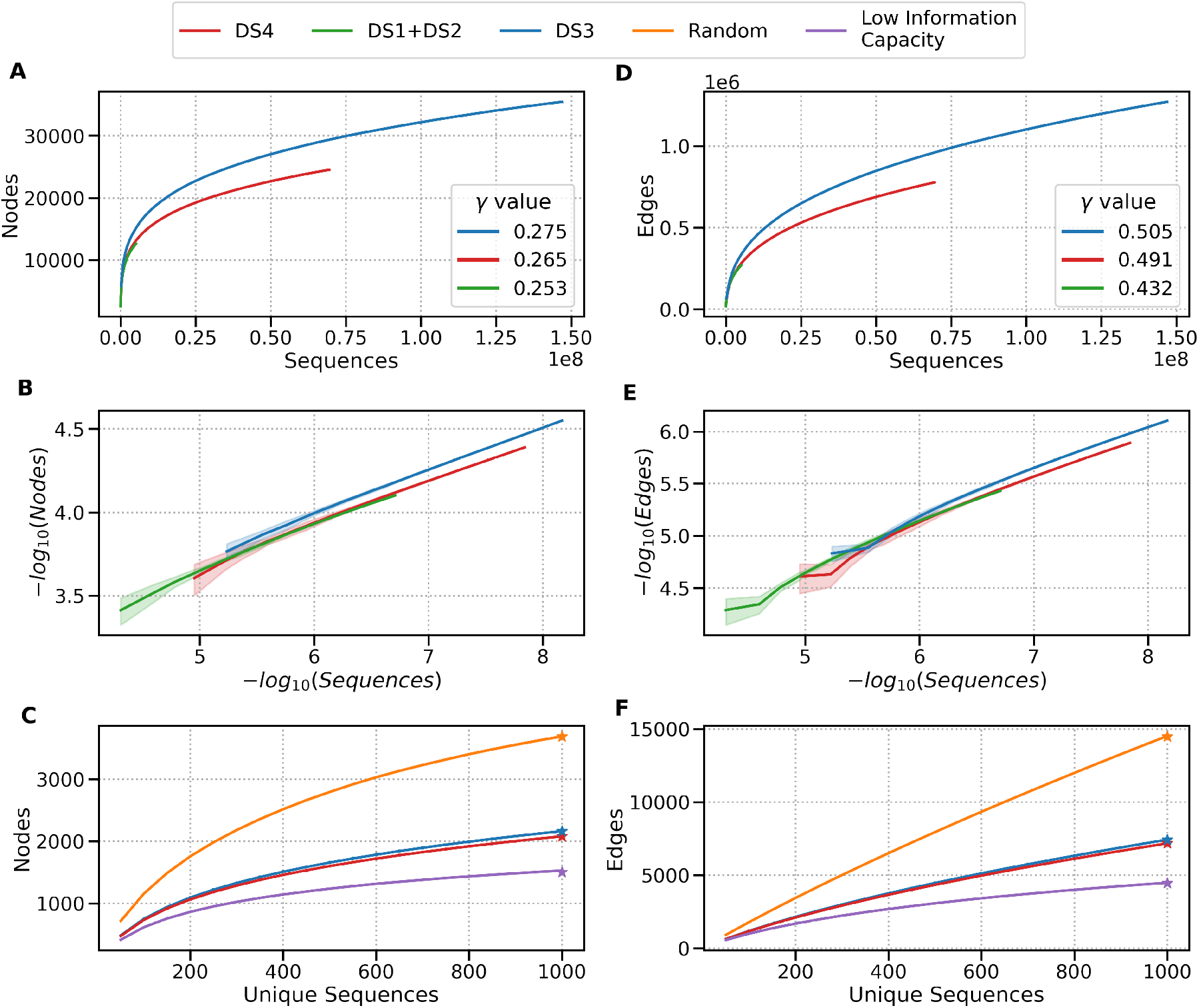
LZGraph Graph Nodes and Edges Growth Rate as a Basis for K1000 Diversity Index. (**A+B**) show the growth in the number of graph nodes based on the number of sequences used in both regular and log-log scales. (**D+E**) shows the growth in the number of graph edges based on the number of sequences used in both regular and log-log scales. (**C+F**) show 4 examples of the K1000 index,the orange curve represents the K1000 index for randomly generated sequences. The blue and red curves represent 30 samples each for DS3 and DS4, respectively. The Purple curve represents the K1000 index for the same number of unique sequences as in all other examples generated using an LZGraph built from only 2000 unique sequences

It is interesting to note that according to Turán’s theorem, the maximum number of edges in a directed graph composed of *N* nodes is *N* ^2^*/*4. As seen in Table 2, indeed the exponent of the nodes is slightly more than one half of the exponent of the edges. However, the moderate differences in the ratios between the exponents of the edges and those of the nodes might indicate non-trivial differences in repertoire diversity measures.

* Where *γ* is the power law coefficient, namely, *f* (*x*) = *a* · *x*^*γ*^

#### LZGraph Based Repertoire Diversity Index

A fundamental task in AIRR-seq analysis involves diversity estimation. Commonly, one first assigns sequences into clones, and then estimates the clonal abundance distribution with a measure of alpha diversity. These measures are borrowed from ecology and include (species) richness, Simpson’s index, Shannon’s entropy, or a continuum of measures suggested by Hill [12]. As such, these measures characterize the distribution of reads attributed to each unique sequence, not taking into account the (dis)similarity between the sequences in a repertoire. Here, we harness the LZGraph data structure and suggest a new measure of repertoire diversity, which goes beyond the clonal abundance distribution. Specifically, we start with the set of unique sequences (which are identical to clones in T cell repertoires) present in a repertoire. We then repeatedly sample this set (without replacement) and construct an LZGraph from each sample. This process is visualized in Figure 5, and new diversity measures can be defined from it. We show here the use of K1000, defined as the number of nodes in an LZGraph constructed from 1000 randomly sampled unique sequences averaged over 50 sampling procedures. As seen in Figure 5, the differences between DS3 and DS4 are maintained in a similar fashion to what we saw in Figure 5. Moreover, the empirical K1000 values of DS3 and DS4 are lower than totally random sequences but higher than those of sequences that were generated from an LZGraph built from 2000 unique sequences only.

Since K1000 is calculated from randomly drawn unique sequences, it is not influenced by the clonal abundance distribution. K1000 values are indicative of the information capacity of the corresponding LZGraph, and as such reflect another type of diversity information. For example, two repertoires can have the same number of unique sequences and the distribution of frequency over these unique sequences. This will result in similar values of alpha diversity but with different K1000 values (“information capacity”, see Figure 5).

#### LZGraph Based Feature Vectors Are Robust Features For Classification Tasks

Similar to the experiment we conducted with the LZ-76 BOW feature vector, we tested the capability of the eigen-centrality values of a Naive LZGraph to function as a feature vector by stratifying females and males in four distinct datasets. Here as well, we observed high mean F1 score on 15 folds of cross-validation and robustness when introducing an unseen dataset (Figure 4A).

#### Assessing Generation Probabilities with LZGraphs

An LZGraph stores the information about the relative probabilities to go from one node to a following node that is directly connected to it. These probabilities can yield a generation probability for each sequence (*P*_*gen*_), by multiplying the values attributed to each edge along the path associated to the sequence. To validate the *P*_*gen*_s generated via the “Amino Acid Positional” LZGraph, we compared the *P*_*gen*_s produced by it to *P*_*gen*_s produced by “Sonia’s” LeftPosRightPos model [32]. For this, we calculated both LZGraph’s and Sonia’s “*P*_*gen*_s” for each sequence in a randomly chosen repertoire from DS1. We observed a high correlation between the two *P*_*gen*_ sets (See Figure 6A). The deviations between the two seem to stem from Sonia’s tendency to attribute extremely low *P*_*gen*_ values in some cases.

**Figure 6:**
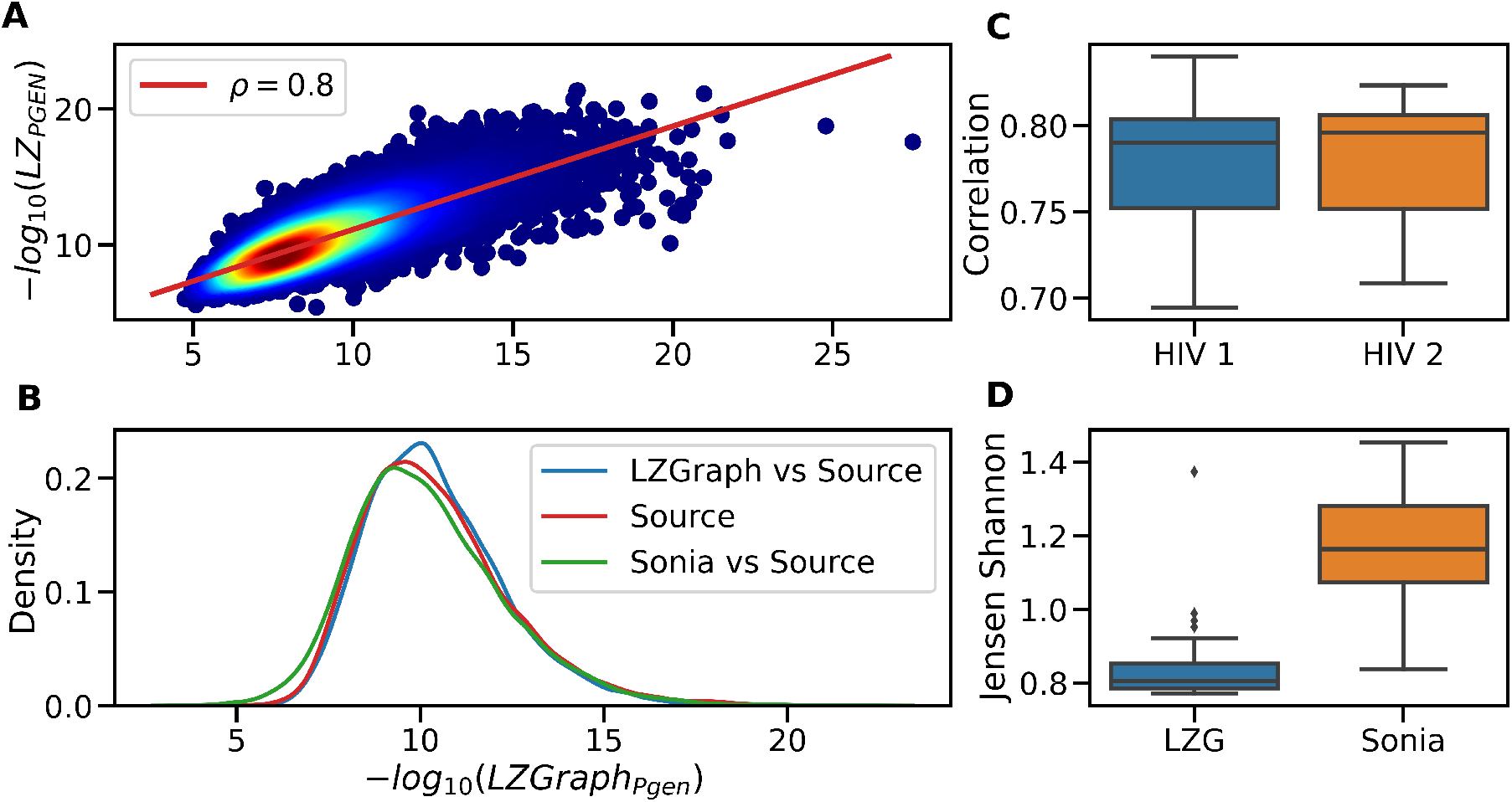
Assessment of LZGraph Generated Sequences and *P*_*gen*_ Quality. (**A**) Shows the correlation between the -log10 *P*_*gen*_ given by SoniaLeftposRightpos and the “Amino Acid Positional” LZGraph for an arbitrary repertoire from DS1. The red line represents the linear fit with Pearson’s correlation equals to 0.8 and slope equals to 0.76. (**B**) Shows the distribution of the LZGraph *P*_*gen*_ values between sequences generated via SoniaLeftposRightpos model after fitting it to the source repertoire and sequence generated using the LZGraph in comparison to the source repertoire. (**C** The Pearson’s correlation between all sample *P*_*gen*_s in both DS1 and DS2 calculated using Sonia and LZGraph, respectively. (**D**) For all samples in DS2, an LZGraph and a SoniaLeftposRightpos model were constructed. The same number of sequences was generated for each repertoire. The LZGraph *P*_*gen*_ was calculated for sequences generated by both models. The distribution of *P*_*gen*_ values in both sets of generated sequences was compared to the distribution of *P*_*gen*_ values derived from the source repertoire. The distribution of the Jensen–Shannon divergence between generated and source sequences is shown for each model separately.

It is important to note that in contrast to Sonia, *LZ* −*P*_*gen*_ does not require gene annotations and is only based on the sequential relationships between LZ-76 sub-patterns in the graph.

#### In-silico Generation of Sequences Using an LZGraph

In addition to estimating generation probabilities with an LZGraph, we can leverage the same information stored in the graph to generate artificial sequences. The simulated sequences are expected to have identical statistical properties as the source repertoire used to construct the LZGraph. To evaluate the quality of the sequences generated using the “Amino Acid Positional” LZGraph (in the “Genetic Constrained” mode; see methods), we selected an arbitrary repertoire from DS1 and constructed an LZGraph corresponding to that repertoire. Independently, we fitted a Sonia model to the same repertoire, and then generated N sequences from both models, where N is the depth of the original repertoire (Figure 6A). LZGraph *P*_*gen*_s were calculated for the source repertoire and the two sets of generated sequences, as can be seen in Figure 6B. When assessing the Jensen-Shannon distance between the generated sequences and the source repertoire, we observe that the LZGraph generated data is closer to the source repertoire in terms of LZGraph generation probabilities. We repeated the same experiment for all repertoires in DS2, and saw identical trends of high correlation between Sonia *P*_*gen*_ and LZGraph *P*_*gen*_ (Figure 6C), with sequences generated using LZGraph having a lower Jensen-Shannon distance from the source repertoire as apposed to the Sonia generated sequences (Figure 6D).

In an opposite setting, when measuring the Jensen-Shannon distance from the source repertoire via the Sonia post *P*_*gen*_, Sonia is closer to the source repertoire than the LZGraph generated sequences (see supplementary figure-3). In summary, both methods to generate artificial sequences and to estimate generation probabilities (LZGraph and Sonia) are consistent when assessing the generation probabilities of the sequences generated by the same method. On the other hand, there is a slight difference when assessing generation probabilities of sequences generated by the other method. The main advantage of the suggested approach (LZGraph) is that there is no need for further annotations to calculate *P*_*gen*_s and to generate artificial sequences that follow the statistical properties of a given repertoire.

### Repertoire-specific LZGraph can be used for advanced analyses of individual sequences

The presented approach reveals a plethora of new ways to explore AIRR-seq data. We show below three examples of new ways to analyze with an LZGraph individual sequence in the context of a specific repertoire.

#### Assessment of Sequence Centrality

An LZGraph is composed of regions of varying densities. Regions in the graph with higher densities have larger support from the sequences used to construct the graph. The density of a region can be used to assess the path that corresponds to a given sequence. As an example, we took two sequences from an arbitrary repertoire from DS1. An LZGraph was constructed from all sequences in the given repertoire, and two sequences were chosen according to their mean Levenshtein distance [23] averaged over all other sequences. The first sequence (“common”) has a much lower value (8) than the other sequence (“rare”), indicating that the “common” sequence is closer on average to the rest of the repertoire compared with the rare one. We then plotted the number of possible alternatives (edges) at each sub-pattern (node) along the sequence for both sequences (Figure 7A). The rare sequence, as calculated by the mean Levenshtein distance to all other sequences, has a lower number of edges along the sequence compared to the common one, and the rate at which the curve approaches zero is also faster for the rare sequence. To generalize this observation, we calculated for each sequence in all repertoires of DS1 these two measures, i.e., mean Levenshtein distance and mean number of edges along the sequence, and observed high Pearson’s correlations between them (−0.63 ± 0.015; supplementary figure-2). Note that at each position of the input sequence, if that position shares the same sub-pattern, it will have the same value in the curve, thus allowing us to assess the degree of difference by quantifying the difference between two or more such curves.

**Figure 7:**
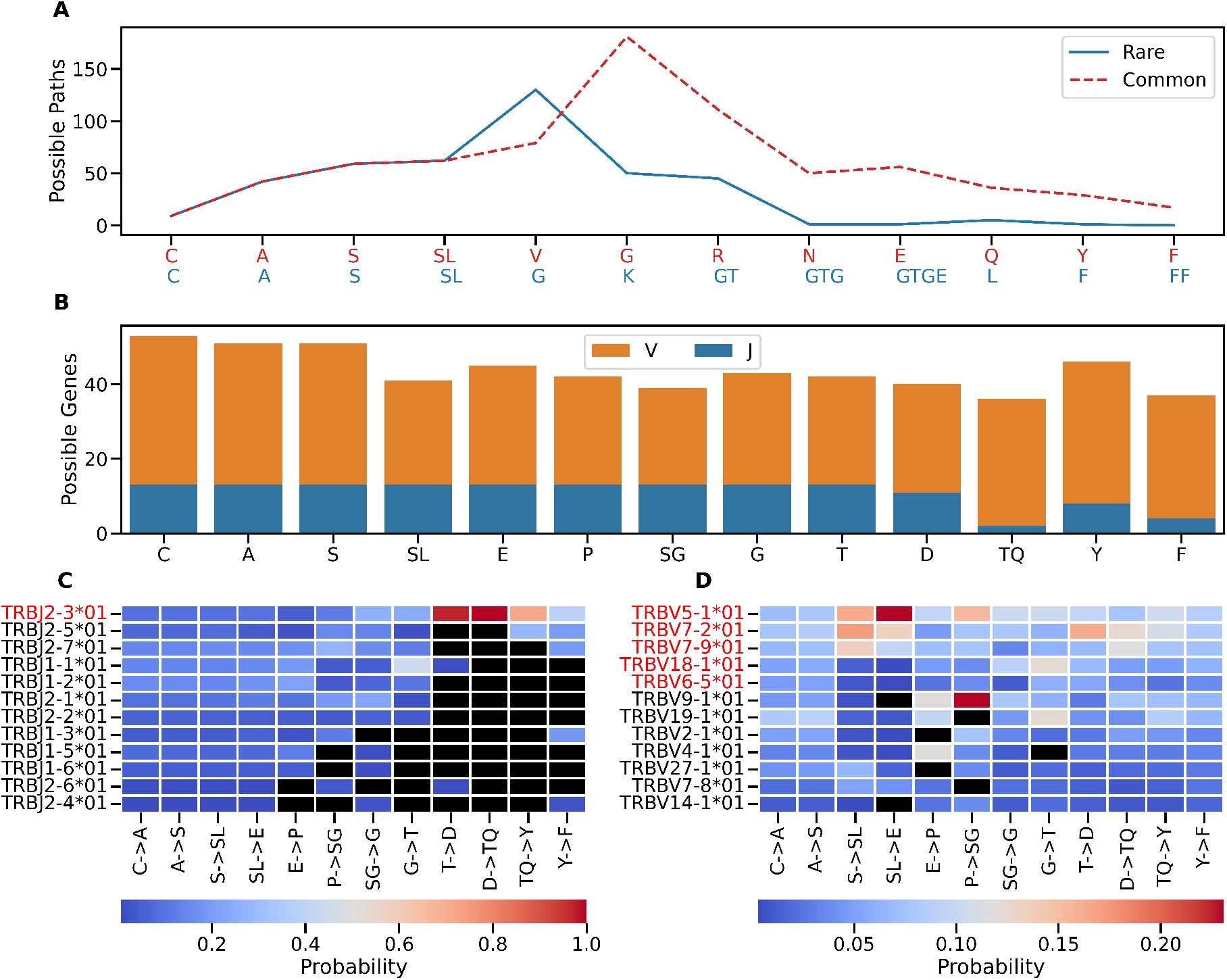
Examples of Analysis Methods for Individual Sequences Using an LZGraph. **(A)** Shows a curve for two sequences where the value of the curve at each sub-pattern is the number of immediately reachable nodes in an LZGraph. A rare and common sequence in blue and dashed red, respectively. (**B**) The number of unique V and J genes that could be used at each sub-pattern position. (**C+D**) The heat maps show all the unique V and J alleles associated with each edge of sequence. The color scale at each cell represents the probability of choosing the gene at that edge. A black cube represents that an allele was never observed at that edge, and a red-colored allele name means that it is observed throughout the entire sequence path.

Not only does the information retrieved from such a representation correlate with the mean Levenshtein distance to all other sequences, but it is also indicative of the centrality of the sub-patterns present in the sequence. It is important to note that the suggested approach is much more efficient in terms of run-time and space complexity. Deriving the analysis presented in Figure 7A for all sequences in a repertoire costs 𝒪 (*N* · *L*) to construct the graph, where *N* is the number of sequences in the graph and *L* is the typical sequence length. This is followed by 𝒪 (*L* · *N*) for the calculation of the possible paths of all sequences, resulting in a total inference time of 𝒪 (2*N* · *L*). On the other hand, calculating the pairwise Levenshtein distance costs 𝒪 (*N* ^2^ · *L*^2^).

#### An LZGraph can store information about flanking genomic regions

An important piece of information when analyzing TCR sequencing data is the identity of the regions that flank the CDR3; namely, the TRBV and TRBJ alleles. This information can be incorporated in an LZGraph by recording the TRBV and TRBJ allele annotations along the path of each sequence that was used to construct the graph. The resulting LZGraph can be used to gain insights on the relationships between sub-patterns (nodes) and transitions (edges), and the flanking TRBV and TRBJ alleles.

Figure 7B shows an example of what can be learned from such annotated LZGraph on a particular sequence (taken unbiasedly from DS1). As almost all TCRB CDR3 aa sequences start with “CAS”, we see that these initial patterns can indeed be attributed to all possible TRBV and TRBJ alleles. On the contrary, the “TQ” sub-pattern of this sequence for example, has only two possible J alleles and 36 possible V alleles. Such a representation allows us to distinguish between nodes that can appear in all/many flanking alleles and nodes that are allele-specific, from which allele annotations can be inferred.

To gain more information about the sequence shown in Figure 7B, we applied an analogous logic to the edges as well. From Figure 7C we can see that only a single TRBJ allele (TRBJ2-3*01) is consistent with the given sequence, mainly due to two edges with one possible allele (*T* →*D*;*D* →*TQ*), and from Figure 7D we see that five alleles have a non-zero probability to be associated with this CDR3.

### LZGraphs Performance Profile

The importance of algorithmic efficiency in large-scale models such as those presented above is crucial. The methods and data structures presented in this paper were implemented using Python 3.9, and performance profiling was carried out to provide the reader with a summary of the computation time scales for the methods. On top of the fact that the methods presented here do not require a costly preliminary annotation step, they are highly efficient in terms of running times (Table 2).

## Discussion

We explore here a new graph-based methodology to analyze TCR repertoires, leveraging the LZ-76 compression. Our model provides the user with a framework for an entire repertoire and single sequences’ analysis in an annotation-free setting. Most of the results presented here do not require germline information such as TRBV/TRBD/TRBJ annotations. Nevertheless, the model can incorporate these annotations, and use them for applications such as synthetic sequence generation and genomic inferences. Furthermore, we introduce a novel diversity index (K1000) that aims to quantify structural patterns, providing a proxy for the potential “information capacity” based on aa or nucleotide sequences in an individual’s repertoire. K1000 is conceptually different from common classic diversity measures used today, as it does not rely on unique sequence abundance (see Figure-5), but rather reflects the “information capacity” of sequences in a given repertoire. It opens the door for new ways to assess repertoire diversity and new comparisons between repertoires.

Throughout the paper, we mainly used an LZGraph to represent the statistical features of a single repertoire. However, it can be used to represent more than one individual. For example, LZGraphs can be constructed for all individuals associated with a particular characteristic (e.g., sex, age, ethnicity, or a particular disease). These resulting graphs can be used to identify features that differentiate between the cohorts and may be harder to trace at the individual level. Moreover, such “master-graphs” can be used to infer *P*_*gen*_’s for empirical sequences and to simulate synthetic ones that will better follow the statistical properties of the common denominator of the source cohorts.

There are a few limitations arising from the method due to its specificity to the source repertoire. Naturally, when we construct a probabilistic model based on a single point of reference without any prior, we are limited to the knowledge obtained solely from our observations; thus, two graphs representing two different repertoires will differ in both the nodes and edges they contain. Naturally, the deeper the repertoire used for the construction of an LZGraph, the more confident the user can be of any difference between two repertoires originating in actual biological differences rather than an effect due to under-sampling of the repertoire. Creating a “master-graph” can overcome the depth issue and assess the difference between “master-graphs” rather than between individual repertoires.

The methodology proposed in this paper uncovers an untapped source of feature extraction based on data and string compression algorithms. Although we used the classic LZ-76 algorithm as the compression algorithm in this paper, it is of high interest to explore other compression algorithms for this purpose in future research. In addition to different compression algorithms, it is interesting to explore whether the genomic information stored in the graph stores information that can be used to infer the germline alleles adjacent to the CDR3 region, namely, the TRBV and TRBJ alleles. While the results presented here are based on TCR CDR3 regions, an analogous methodology can be applied to other types of sequences as well. Specifically, it will be interesting to adapt the presented method to BCRs, and explore how somatic hypermutations affect the analysis results. It might even reveal new insights about the somatic hypermutation mechanism.

In summary, this paper took the first step in representing TCR repertoire data using lossless compression algorithms. Future work is needed to explore the many possible extensions and adaptations of the presented approach in the context of AIRR-seq and other types of biological sequences.

## Competing interests

The authors declare that they have no competing interests.

## Code Availability

All the described above methods are implemented and packaged into a convenient plug and play python package, For further references see https://github.com/MuteJester/LZGraphs

## Acknowledgements

This study was partially supported by grants from the ISF (2940/21), VATAT grant, and the European Union’s Horizon 2020 research and innovation program (825821 to GY). The contents of this document are the sole responsibility of the iReceptor Plus Consortium and can under no circumstances be regarded as reflecting the position of the European Union.

## Notes

https://github.com/MuteJester/LZGraphs

### Competing Interest Statement

The authors have declared no competing interest.

