## Supplementary material for "A Novel Approach to T-Cell Receptor Beta Chain (TCRB) Repertoire Encoding Using Lossless String Compression"

Thomas Konstantinovsky Gur Yaari

January 30, 2023

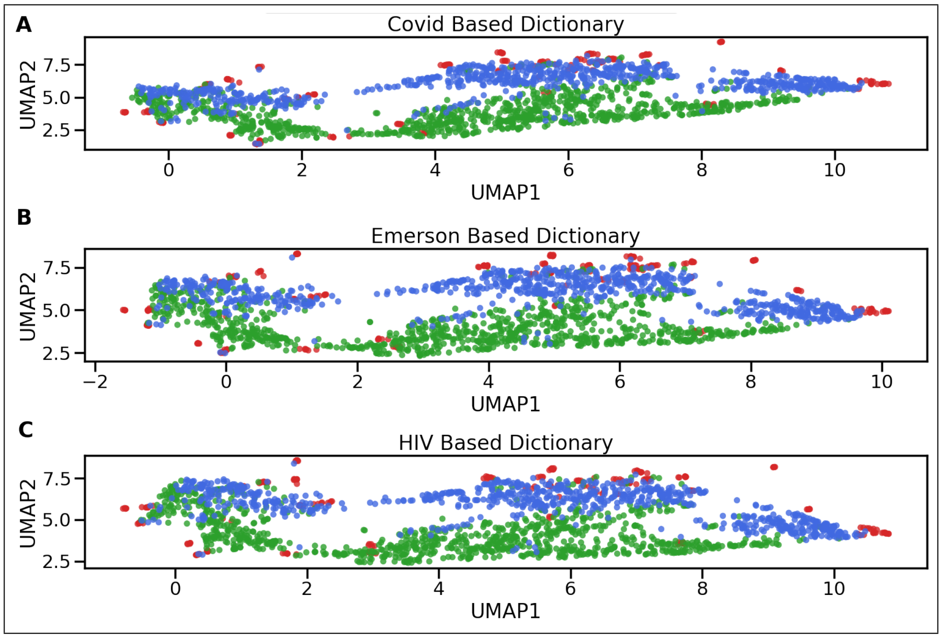


Figure 1: Figures show the projection onto $R^2$ using UMAP of the datasets used in the paper. At each panel the datasets were encoded using a BOW dictionary derived from only one dataset. The dictionaries in panels (A-C) are DS1+DS2,DS3 and DS4, respectively.

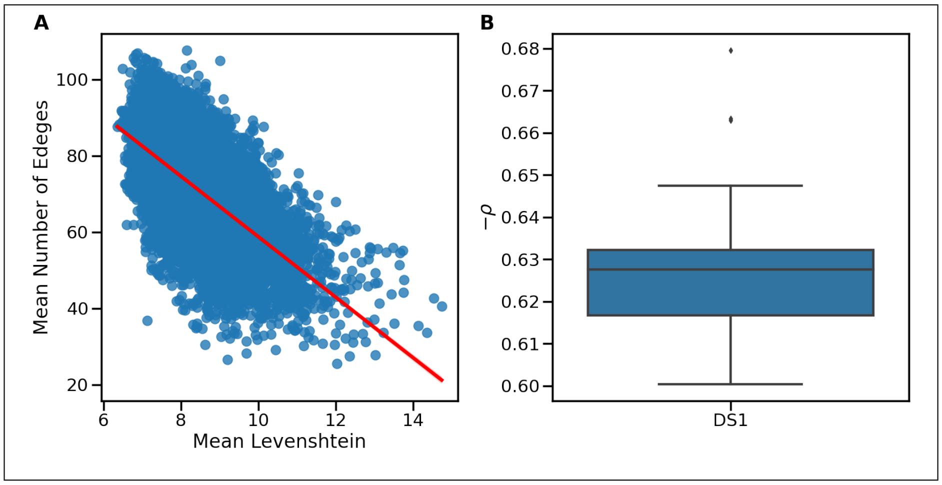


Figure 2: (A): the mean Levenshtein distance between all the sequences in an arbitrary repertoire (x-axis) against the mean number of edges throughout the path in the corresponding LZGraph (y-axis). (B): the correlation between the mean Levenshtein distance between all the sequences and the mean number of alternative edges throughout the path in an LZGraph, calculated on all repertoires in DS1

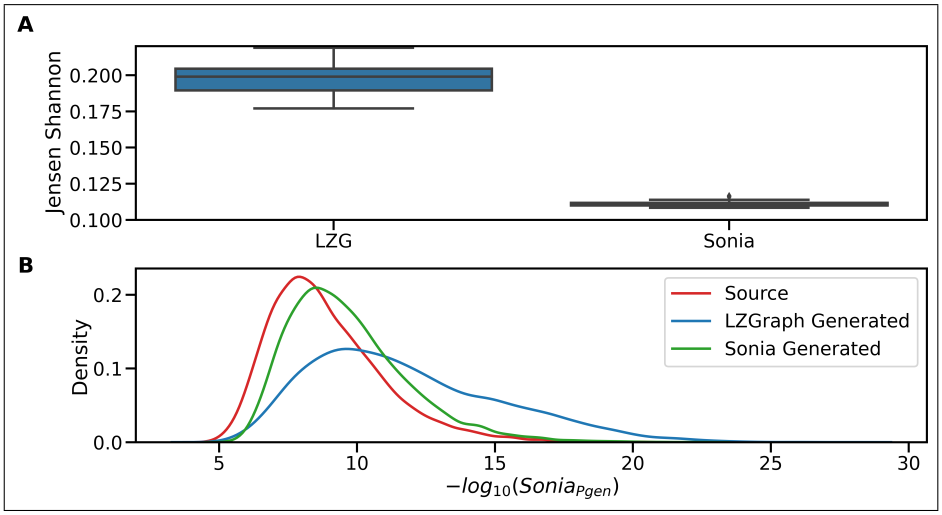


Figure 3: (A): the Jensen Shannon distance between the Sonia Post Pgen between the source repertoire used to construct an LZGraph and Sonia LeftPosRightPos model to sequence generated from both models.(B): an arbitrary sample from DS1 was taken, example of Sonia Post Pgen on the source sequences, generated using Sonia sequences and LZGraph generated sequences.
